# Tree Explorer (T-REX): Bridging Phylogenetic and Phenotypic Data for Enhanced Analysis and Interpretation

**DOI:** 10.64898/2025.12.01.691494

**Authors:** Damian Magill, Jean-Marc Ladrière

## Abstract

**Background:** The Tree Explorer (T-REX) project is an open-source initiative designed to enhance the visualization and integration of phylogenetic and phenotypic data. Utilizing the ete3 toolkit, T-REX offers a user-friendly interface and a suite of tools written in Python. This project addresses the longstanding need to easily label phylogenetic trees with phenotypic data stored in other files and efficiently query this data to find entities of interest.

**Results:** T-REX includes various tools to help users prepare their input files and refine the output to create publication-worthy figures. It aims to facilitate the rapid identification of strains of interest for research projects, making large phylogenetic trees more interpretable and useful for evolutionary biology studies.

**Conclusions:** T-REX provides an easy-to-use tool that allows users to effectively link their phylogenetic and phenotypic datasets in order to answer diverse biological questions. Available as a pre-compiled binary or rapid install, users can label trees, perform complex queries to identify individuals of interest, link large-scale quantitative datasets, and produce publication-ready figures. This tool, along with its future developments, is expected to assist in evolutionary biological studies and in the identification and selection of biological entities exhibiting traits of interest.

## Introduction

The reconstruction of phylogenetic relationships between living organisms is a cornerstone of evolutionary biology. Understanding these relationships offers unparalleled insights into the phenotypic characteristics of organisms through ancestral inference. By combining phenotypic data with in-depth genomic analyses, researchers can trace the loss and gain of traits and identify processes like convergent evolution. Initially, phylogenetic relationships were constructed using morphological characteristics, but the advent of first-generation sequencing technologies shifted the focus to molecular methods. With the rise of high-throughput methodologies, we are transitioning from the phylogenetic era to the phylogenomic era and enhancing our ability to generate data on various phenotypic characteristics. Despite this explosion of data, challenges exist at all levels with respect to the accurate reconstruction of phylogenetic relationships and their exploitation [1].

Numerous tools exist for inferring phylogenetic relationships and visualizing trees. Popular packages include RAxML, MrBayes, and IQ-TREE for relationship inference [2]; [3]; [4] while MEGA and Geneious offer both inference and tree visualization capabilities [5]. Visualization tools like FigTree and Interactive Tree of Life (IToL) are also crucial for rendering large phylogenetic trees interpretable and providing a means to produce high quality publication ready figures [6]. Despite these tools, there remains a gap in effectively linking phylogenetic data with complex phenotypic information. The ability to easily associate traits with evolutionary relationships would enable rapid identification of individuals sharing characteristics, which is of significant interest. The Tree Explorer (T-REX) project is an ongoing open-source initiative leveraging the ete3 toolkit [7]. It provides a user-friendly interface and a suite of tools written in Python for visualizing and integrating phylogenetic and phenotypic data.

## Implementation

The T-REX tool suite has been developed entirely in Python, utilizing the ete3 toolkit as a foundational framework for manipulating tree objects [7].

T-REX can be executed directly from the source code, following the installation of a few dependencies, or it can be installed via pip. Additionally, standalone executable binaries for Windows are available through the GitHub repository, providing a convenient installation option. The package has been created with users in mind with error and information messages designed to be informative.

Comprehensive instructions are available at https://github.com/DamianJM/T-REX/.

The test files used to demonstrate the functionality of T-REX are also accessible through the public repository.

## Results and Discussion

### Phenotype Mapping and Querying

To conduct a phenotype mapping-associated analysis, begin by preparing the traits file. Ensure that the first column contains a series of identifiers that correspond to the names used in the phylogenetic tree. The file can be in Excel or CSV format, but it is advisable to avoid using an excessive number of special characters.

To facilitate the creation of compatible files, several tools have been provided, notably the “Tree Name Export” function, which rapidly extracts names from phylogenetic trees to aid in downstream trait file creation. This is particularly useful when the user starts with only a tree file.

Once the file is prepared, upload it and click on “label options” to select the features of interest. By default, no features are selected to prevent overcrowding of the result output. After making your selections, upload the tree, and the selected labels will appear beside the corresponding strain matches. Absent identifiers will be ignored.

Next, utilize the powerful label search feature to query the tree. Enter your search criteria in the query box using the trait file label names and either absolute values or ranges, depending on the data type. For example, to highlight strains that do not contain the gene g1 and have an average colony size under 7 mm, use the query “g1=0 AND colony size=(L, 7) AND colour=yellow” (see Figure 1). Queries can be as complex as necessary and can also be used to pre-collapse or crop sections of the tree to simplify the view, especially for large and complex trees.

**Figure 1.**
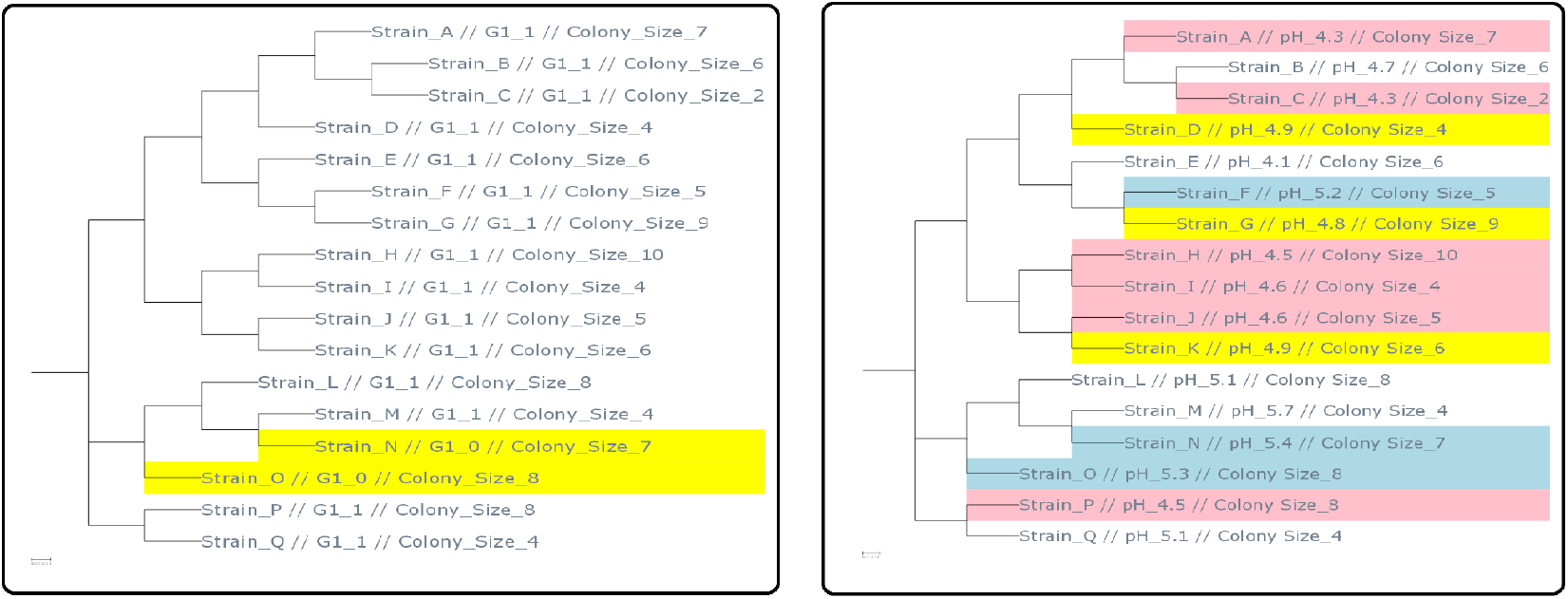
Example T-REX output on test data following specific query for g1 gene absence and colony size less than 7mm (left) and advanced multi-query highlighting different final pH values and colony sizes (right)

Multiple queries can be written that will be executed sequentially, providing a potentially powerful means to identify strains of interest.

### Quantitative Data Displays

When handling arrays of quantitative data, users can perform complex queries to select strains based on various ranges and thresholds. However, this approach may not always be practical. A more effective method is to use a visual display of the data. This is provided as a distinct heatmap functionality within the application, allowing users to easily link and display data directly on the tree. This feature is particularly useful for visualizing quantitative data from large-scale phenotypic experiments and core-pan genomic datasets, where users can utilize presence-absence data or specific numeric labels corresponding to gene variants Figure 2.

**Figure 2.**
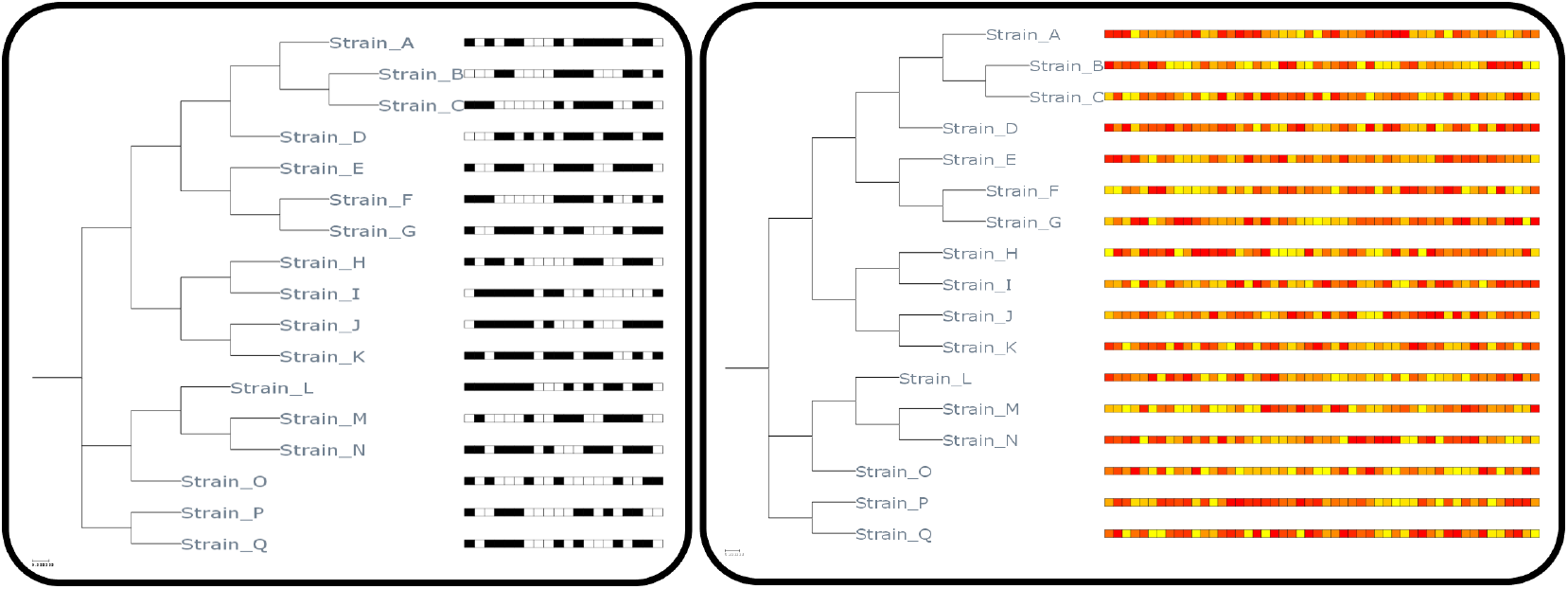
Example output using heatmap functionality on presence/absence data (left) and quantitative data (right) with different formatting applied

Once users have displayed their data and applied queries, they can output directly to file or visualise the tree directly and apply additional formatting as necessary.

## Conclusion

T-REX provides an easy-to-use tool that allows users to effectively link their phylogenetic and phenotypic datasets in order to answer diverse biological questions. Available as a pre-compiled binary or rapid install, users can label trees, perform complex queries to identify individuals of interest, link large-scale quantitative datasets, and produce publication-ready figures. This tool, along with its future developments, is expected to assist in evolutionary biological studies and in the identification and selection of biological entities exhibiting traits of interest.

## Declarations

### Ethics approval and consent to participate

Not applicable.

### Consent for publication

All authors agree to the publication of the manuscript in its present form.

## Availability of data and materials

The datasets used and/or analyzed during the current study are available from the corresponding author on reasonable request. Comprehensive instructions and test files demonstrating the functionality of T-REX are accessible through the public repository at https://github.com/DamianJM/T-REX/.

## Authors’ contributions

Damian Magill and Jean-Marc Ladrière conceived and designed the study. Damian Magill developed the T-REX tool suite and wrote the manuscript. Jean-Marc Ladrière contributed to the manuscript revision. Both authors read and approved the final manuscript.

## Acknowledgements

We would like to thank the IFF R&D Culture Development team for their support and valuable discussions throughout the development of the T-REX project. notably Dr Armelle Blachère for valuable feedback.

## Authors’ information

Damian Magill and Jean-Marc Ladrière are researchers at IFF R&D Culture Development, 86220, Dangé-Saint-Romain, France.

## Availability and requirements

- Project name: T-REX
- Project home page: https://github.com/DamianJM/T-REX
- Operating system(s): Platform independent, binary for Windows
- Programming language: Python
- Other requirements:
- License: OPEN MIT
- Any restrictions to use by non-academics: None

## References

[1] P. Kapli, Z. Yang, and M. J. Telford, “Phylogenetic tree building in the genomic age,” Nature Reviews Genetics, vol. 21, no. 7, pp. 428–444, 2020, doi: 10.1038/s41576-020-0233-0.

[2] A. Stamatakis, “RAxML version 8: a tool for phylogenetic analysis and post-analysis of large phylogenies,” Bioinformatics, vol. 30, no. 9, pp. 1312–1313, 2014, doi: 10.1093/bioinformatics/btu033.

[3] J. P. Huelsenbeck and F. Ronquist, “MRBAYES: Bayesian inference of phylogenetic trees,”Bioinformatics, vol. 17, no. 8, pp. 754–755, 2001, doi: 10.1093/bioinformatics/17.8.754.

[4] L.-T. Nguyen, H. A. Schmidt, A. Von Haeseler, and B. Q. Minh, “IQ-TREE: a fast and effective stochastic algorithm for estimating maximum-likelihood phylogenies,” Molecular Biology and Evolution, vol. 32, no. 1, pp. 268–274, 2015, doi: 10.1093/molbev/msu300.

[5] K. Tamura, G. Stecher, and S. Kumar, “MEGA11: molecular evolutionary genetics analysis version 11,” Molecular Biology and Evolution, vol. 38, no. 7, pp. 3022–3027, 2021, doi: 10.1093/molbev/msab120.

[6] I. Letunic and P. Bork, “Interactive Tree Of Life (iTOL) v5: an online tool for phylogenetic tree display and annotation,” Nucleic Acids Research, vol. 49, no. W1, pp. W293–W296, 2021, doi: 10.1093/nar/gkab301.

[7] J. Huerta-Cepas, F. Serra, and P. Bork, “ETE 3: reconstruction, analysis, and visualization of phylogenomic data,” Molecular Biology and Evolution, vol. 33, no. 6, pp. 1635–1638, 2016, doi: 10.1093/molbev/msw046.

